# Potential soil transmission of a novel *Candidatus Liberibacter* strain detected in citrus seedlings grown in soil from a huanglongbing infested citrus grove

**DOI:** 10.1101/821553

**Authors:** Ulisses Nunes da Rocha, Keumchul Shin, Sujan Timilsina, Jeffrey B. Jones, Burton H. Singer, Ariena H. C. Van Bruggen

## Abstract

*Candidatus* Liberibacter spp. are *Alphaproteobacteria* associated with plants and psyllid vectors. Most cause plant diseases, including *Ca* Liberibacter asiaticus (Las) associated with citrus huanglongbing (HLB). Replacing HLB-infected by Las-free citrus trees results in fast re-infection despite psyllid control. To check if HLB could be soil-borne, we performed an insect-free greenhouse-experiment with 130 mandarin seedlings in two citrus-grove soils (A and B), non-autoclaved or autoclaved. Liberibacter-specific 16S-rDNA PCR primers to detect Las were used to search for *Ca.* Liberibacter spp. in mandarin leaves. Seven plants grown in non-autoclaved soil B showed HLB-like symptoms and tested positive after 2.5 and 8.5 months using three different primer systems: two based on the 16S-rDNA gene (primers HLBas/HLBr and OI2c/OI1) and one based on the rplA/rplJ gene (primers LAA2/LAJ5). DNA segments from these plants amplified by primers OI2c/OI1 were cloned and sequenced; they were 95.9 % similar to Las and 94.8% to *Ca.* Liberibacter africanus (Laf). The DNA product from Liberibacter-group specific PCR primers for the rplA/rplJ gene was 87.6% similar to that of Las and 78.2% of Laf. As the strain obtained originated from soil and was different from existing *Ca*. Liberibacter species, this strain may be a new species.

## INTRODUCTION

*Candidatus* Liberibacter is a group of *Alphaproteobacteria* that can cause serious diseases in plants. The name *Candidatus* Liberibacter was proposed in 1994 (Jagoueix et al. 1994) and the first species *Ca.* Liberibacter asiaticus (Las) and *Ca.* Liberibacter africanus (Laf) were defined soon after (Planet et al. 1995). A decade later, a novel species belonging to this group was found in Brazil and named *Ca.* Liberibacter americanus (Lam) (do Carmo Teixeira et al. 2005). Recently, a new strain of *Ca*. Liberibacter in citrus in Colombia was provisionally named *Ca*. Liberibacter caribbeanus (Keremane et al. 2015). All the above mentioned species are associated with citrus greening or Huanglongbing (HLB), transmitted by the psyllid vectors *Diaphorina citri* Kuwayama or *Trioza erytreae* (Del Guercio) (Bové 2006; Chiyaka et al. 2012; Shimwela et al. 2016). In citrus, the pathogens reside primarily in the phloem (Bové 2006) and to a lesser extent in the xylem (Ebert et al. 2018). Currently, huanglongbing is the most destructive citrus disease worldwide (da Graça et al. 2016; Farnsworth et al. 2014; Gottwald 2010; Narouei-Khandan et al. 2016; Shen et al. 2013b).

A three-pronged management approach has been adopted in most citrus production areas affected by HLB: (i) production of clean planting stock in the absence of the psyllid vectors, (ii) vector control by insecticides, and (iii) reducing available inoculum by rogueing of infected symptomatic trees (Bassanezi et al. 2013; Belasque et al. 2010; Bové 2014; Salifu et al. 2012). In addition, treatment of infected trees with antibiotics (Blaustein et al. 2018; Shin et al. 2016; Yang et al. 2016) or plant nutritionals and defense enhancing chemicals (Shen et al. 2013a; Xia et al. 2011; Xu et al. 2013), heat treatment (Hoffman et al. 2013; Yang et al. 2016) and selection of resistant or tolerant rootstocks (Wang et al. 2016) have also been attempted. Despite these efforts at managing the disease, HLB has continued to spread in conducive areas (Narouei-Khandan et al. 2016; Shen et al. 2013b; Shimwela et al. 2018a, b). Moreover, planting of disease-free citrus stock in groves where infected trees were removed has often resulted in reinfection of the newly planted young trees by Las despite intensive vector control (Timmer 2014), reminiscent of replant diseases in other fruit trees (Browne et al. 2018; Mazzola and Manici 2012; Yang et al. 2012). However, *Ca.* Liberibacter spp. have never been found in soil thus far, although very closely related genera are often soil-borne (O’Brien and van Bruggen 1991; Shin and van Bruggen 2018; van Bruggen et al. 1990) and Las was detected in colony mixtures isolated from citrus rhizosphere soil on a low-carbon agar medium (Ascunce et al. in preparation).

In addition to the species of *Ca.* Liberibacter associated with HLB, several other members of this genus are associated with plant diseases (Haapalainen 2014). Psyllid yellowing affecting potato and tomato is associated with *Ca.* Liberibacter psyllaurous (Hansen et al. 2008; McKenzie and Shatters 2009). Potato zebra chip disease and tomato decline are presumably caused by *Ca.* Liberibacter solanacearum (Abad et al. 2009; Liefting et al 2009; Thomas et al. 2018), which is considered synonymous with *Ca.* Liberibacter psyllaurous (Morris et al. 2017). A strain of *Ca.* Liberibacter solanacearum, haplotype C, induces symptoms in carrot but not in potato (Haapalainen et al. 2018a, 2018b). In addition, three different species of *Ca.* Liberibacter were discovered: *Ca.* Liberibacter europaeus in the phloem of seemingly healthy pear trees (Raddadi et al. 2011), *Liberibacter crescens* in the phloem of defoliating mountain papaya in Puerto Rico (Fagen et al. 2014a; Leonard et al. 2012), and *Ca.* Liberibacter brunswickensis in Australian eggplant psyllids (Morris et al. 2017). The potential pathogenic nature of these last three species is unknown. So far, only *Liberibacter crescens* has been isolated and maintained in culture consistently (Fagen et al. 2014a, b; Lai et al. 2016). Temporary isolations have been reported for Las (Davis et al. 2008; Parker et al. 2014; Sechler et al. 2009), but the isolates could not be maintained in culture.

The increasing number of species belonging to *Ca.* Liberibacter suggests that there may be other, closely related species or subspecies in unexpected habitats (Haapalainen 2014; Haapalainen et al., 2018b). Las appeared to be quite diverse based on single-nucleotide polymorphism (SNP) analysis of 16S rRNA, ribosomal protein genes, and other gene sequences extracted from HLB-symptomatic citrus trees (Adkar-Puroshothama et al. 2009; Furuya et al. 2010; Katoh et al. 2012). Variability among isolates of Las has also been shown by the variable number of tandem repeats (VNTRs) (Ghosh et al. 2015; Ma et al. 2014; Matos et al. 2013), *omp*-based PCR-restriction fragment length polymorphism (RFLP) (Bastianel et al. 2005; Hu et al. 2011), multilocus microsatellite analysis (Islam et al. 2012), multilocus simple sequence repeat (SSR) profiles (Katoh et al. 2012), and prophage sequence analysis (Jantasorn et al. 2012; Tomimura et al. 2009; Liu et al. 2011; Puttamuk et al. 2014; Zheng et al. 2017; Zhou et al. 2011). Similarly, high genetic diversity was described for *Ca*. Liberibacter solanacearum by SNP analysis of 16S rRNA and ribosomal protein genes (Hajri et al. 2017), multilocus SSR markers (Haapalainen et al., 2018b; Lin et al. 2012), and whole genome sequencing (Lin et al. 2011; Thompson et al. 2015). The variations in Las and *Ca*. Liberibacter solanacearum strains were mostly associated with large geographic regions (Haapalainen et al., 2018a).

Based on the high infection rate of young citrus trees replacing HLB infected trees despite intensive psyllid control (Timmer 2014), the frequent association of Las with roots (Johnson et al. 2014), and the great variability of *Ca.* Liberibacter at the species and subspecies levels, the authors proposed three working hypotheses for the current study: (i) *Ca.* Liberibacter species can be transmitted to healthy citrus seedlings through soil containing residues of HLB affected mature trees, (ii) the transmitted *Ca.* Liberibacter bacteria can induce typical HLB symptoms, and (iii) *Ca.* Liberibacter strains associated with symptoms in replanted citrus seedlings are related to the species associated with HLB affected mature trees. To test these hypotheses we performed a one-year greenhouse experiment in an insect free environment and planted citrus seedlings in two different soils to verify if soil type may influence the *Ca.* Liberibacter spp. that may be transmitted to citrus seedlings. *Ca.* Liberibacter spp. group-specific real time and regular PCR primers were used to detect samples with presumptive Liberibacter-like sequences. Regular PCR primer products were then cloned and sequenced. These sequences were compared with the latest databases. Phylogenetic analysis demonstrated that a novel strain of *Ca.* Liberibacter sp., related to but distinct from *Ca.* Liberibacter asiaticus, was detected in several citrus plants that were never in contact with psyllids.

## MATERIAL AND METHODS

### Soil samples

Soil was collected from two citrus groves in central Florida: (A) the USDA Citrus Research Station in Winter Haven, Polk County, Florida, and (B) a grove in Windermere, Orange County, Florida. Both groves had Hamlin oranges on Swingle citrumelo rootstocks and were managed in a conventional way. In both groves, the trees showed typical symptoms of HLB and representative samples had tested positive for Las with quantitative PCR in 2009, 2010 and 2011 at the Division of Plant Industry (DPI), Gainesville, Florida.

Soil samples were collected on four sides (N, E, S, W) under the canopy of five HLB-positive trees per grove. All roots with a diameter of 5 mm or less were included in each soil sample. Soil A was a yellow-brown fine sandy loam, and soil B a grey-black sandy loam. Air-dried soil samples were subjected to chemical analysis in the Soil Analysis lab at the University of Florida (UF), Gainesville, FL. The pH, soluble P and K contents were very similar at the two locations, but the organic matter and total N contents were significantly higher in soil B than soil A (Table S1). The samples of each of the ten trees were kept separate and can be considered as five replicates within the two groves.

**Table 1.**
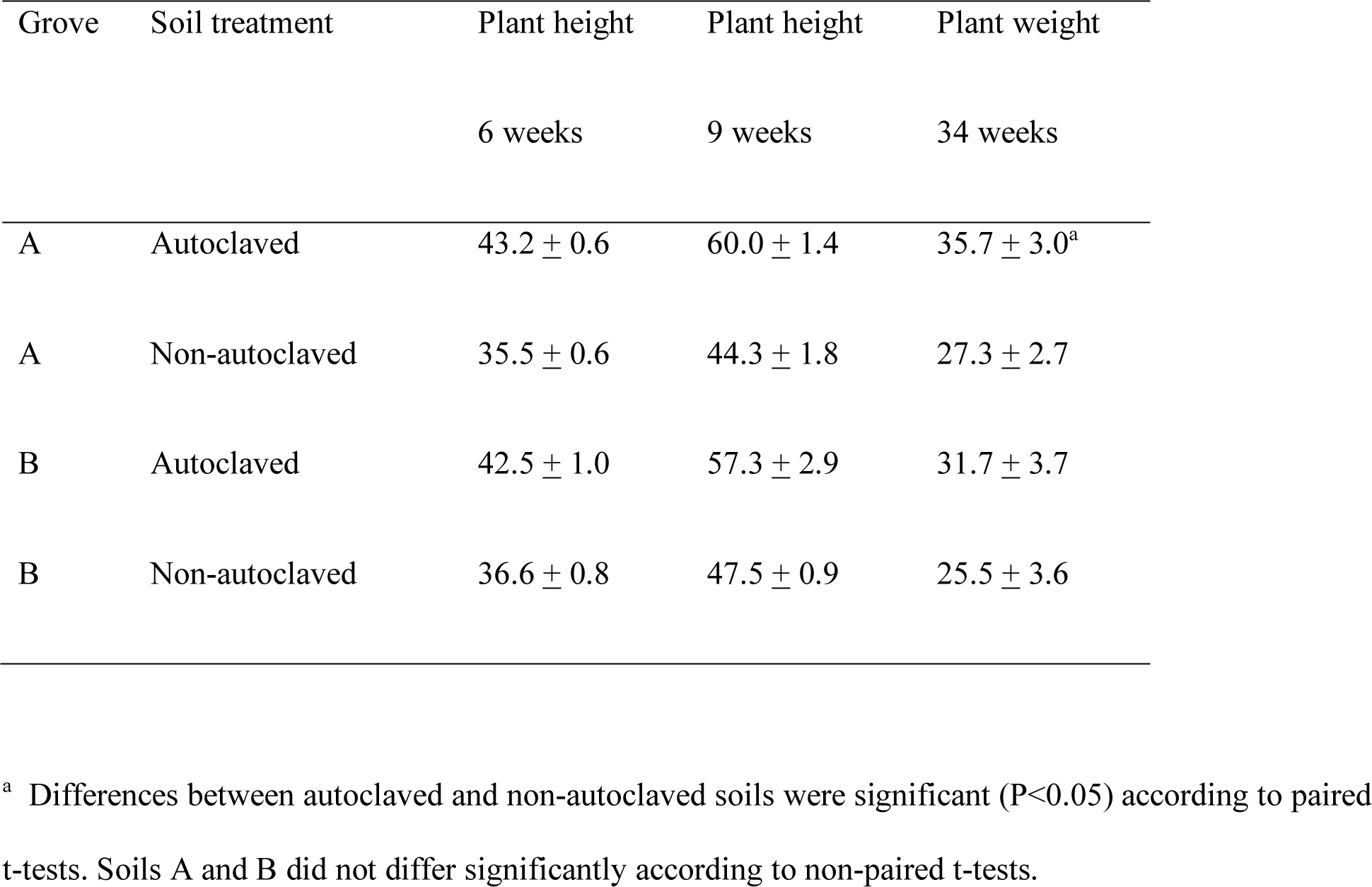
Mean plant heights (cm) and weights (g) and standard errors of the means of mandarin seedlings in autoclaved and non-autoclaved soil from two citrus groves in central Florida after 6, 9, and 34 weeks of growth in a greenhouse in Gainesville, Florida. There were 5 plants per treatment (one for each soil sample around each of 5 trees per grove) in 5 blocks. Differences between autoclaved and non-autoclaved soils were significant (P<0.05).

All soil was sieved through a 1-cm sieve one day after collection. The citrus roots were cut into pieces of 1-2 cm long and returned to the soil. All tools were disinfected with 70% alcohol and all activities were carried out with clean plastic gloves to avoid cross contamination among soil samples. The field capacity of each soil was determined. Field capacities of soil A and B were 20.9% ± 1.7% and 25.5% ± 1.0%, respectively. The moisture contents of the original soil samples were 3.7% ± 0.4% and 8.4% ± 0.7% for soil A and B, respectively.

Half of each soil sample was autoclaved at 120°C in double autoclave bags for 50 min, and left open on a greenhouse bench. In order to promote colonization of the autoclaved soils by bacteria (to avoid ammonia toxicity and poor plant growth), a suspension of naturally occurring soil bacteria was added, prepared with soil from an organically managed experimental vegetable field in Gainesville. The bacterial density was determined in a spectrophotometer at 630 nm using a standard density curve. After 10-fold dilution, 200 ml of suspension was added to each subsample of 4 liters of autoclaved and cooled soil, resulting in a bacterial concentration of 10^7^ CFU/g of dry soil. The amended soil samples were mixed thoroughly in plastic bags. The microbial community was allowed to grow and equilibrate for two weeks. Details on soil collection, soil analyses, and soil treatments can be found in the supplement.

### Experimental set up and plant management

Five-month old mandarin seedlings ‘Cleopatra’ on their own roots were obtained from a citrus nursery producing certified HLB-free trees (Brite Leaf Nursery LLC, Lake Panasoffkee, Florida). The seedlings were transplanted in the autoclaved and non-autoclaved soil samples in five randomized complete blocks, one seedling per soil sample per block (for a total of 5×5×2×2=100 mandarin trees). Thirty residual seedlings were left in pasteurized potting mix. The pot size was 2 L. The pots with autoclaved and non-autoclaved soil from the same trees were placed side-by-side on greenhouse benches for paired comparisons (Fig. 1). The mandarin plants were watered by drip irrigation to avoid microbial cross contamination between pots. Details of plant maintenance are described in the supplement.

**Fig. 1.**
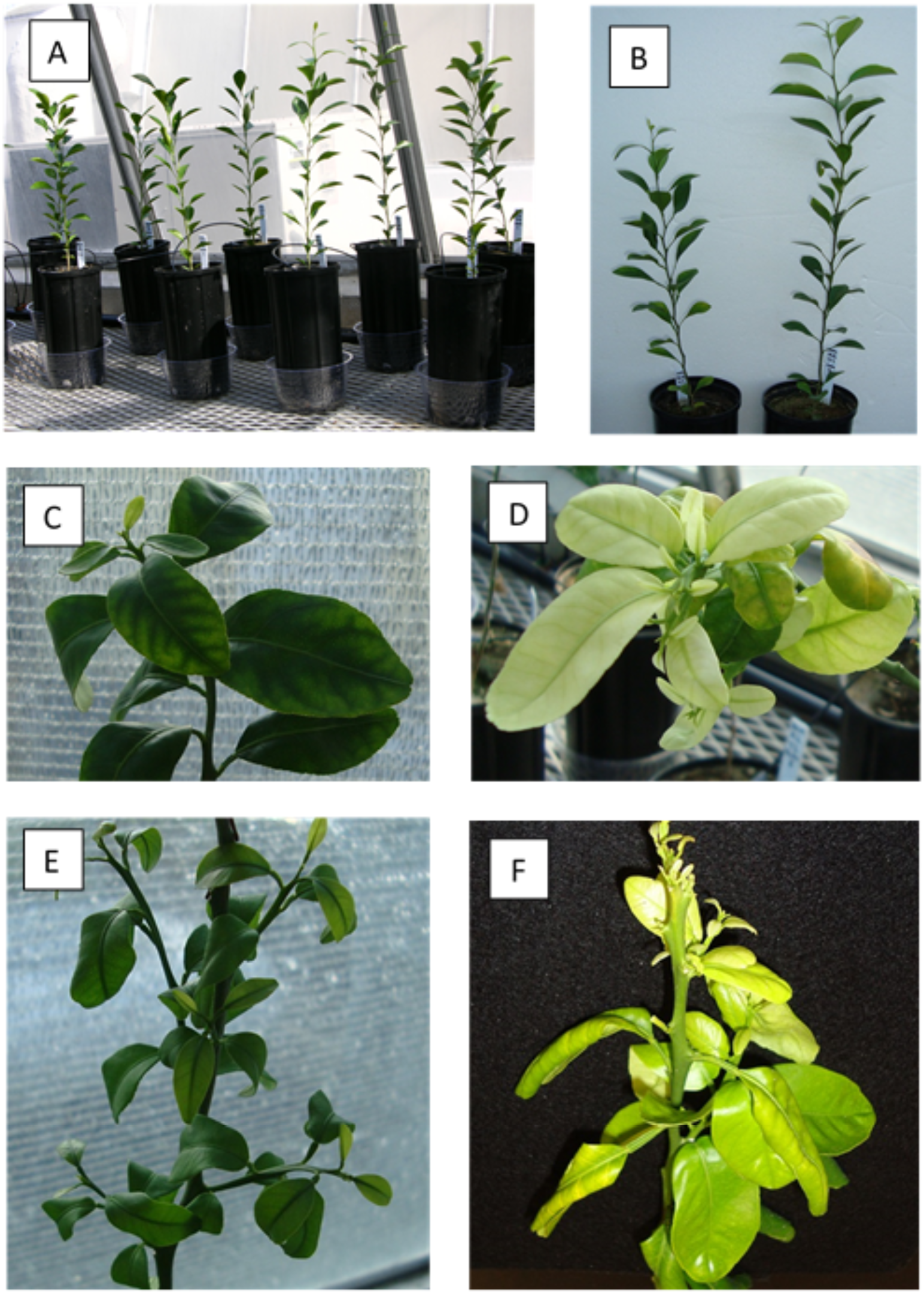
Mandarin seedlings growing in soil collected from two citrus groves in a greenhouse experiment at Gainesville, Florida: (A) experimental setup of seedlings in autoclaved and non-autoclaved soil; (B) typical plant height in non-autoclaved (left) and autoclaved (right) soil; (C) plant in autoclaved soil from grove B; (D) plant in non-autoclaved soil from grove B; (E) plant in autoclaved soil from grove B; (F) plant in non-autoclaved soil from grove B. Pictures (A) and (B) were taken 2 and 2.5 months after planting, pictures (C), (D), (E) and (F) 8.5 months after planting. Plants (D) and (F) were growing in soil R1T9, where earlier samples (2.5 months after planting) were Las-positive in a qPCR test and were related to Las in sequence analyses.

### Plant observations and sample collection

The plants were observed once a week for symptom development. Two young (but completely developed) leaves were harvested from plants with mild discoloration two months after planting using gloves sprayed with 70% alcohol before each new plant, and stored in plastic bags in the refrigerator. Two weeks later, all citrus trees in nonautoclaved soil of blocks 1 and 5, and two trees in autoclaved soil of the same blocks were collected. The shoots were cut off and placed in a plastic bag as described above. The roots were inspected for insects and disease symptoms. Over a period of one week, petioles and midribs of all leaves stored in the refrigerator were dissected out with a sterile scalpel, weighed (107 ± 21 mg per sample), placed in eppendorf tubes and stored in the −80 freezer.

Eight and a half months after planting, the remaining citrus trees were sampled as described above. The shoots were cut off, weighed, placed in a plastic bag, and stored in the refrigerator. The roots were inspected as described above. The petioles and midribs of all leaves were dissected out, weighed, placed in eppendorf tubes and stored in the −80 freezer for DNA extraction later.

Treatment effects (autoclaved versus non-autoclaved soil) on all plant measurements were determined by paired t-tests in Microsoft Excel, as the experiment had a randomized complete block design with soil source (grove and tree number) in main ‘plots’ and soil treatment in ‘subplots’. The groves were compared by non-paired t-tests considering the five trees as replicates.

### DNA Extraction

Midribs of top leaves of each plant were cut (approximately 1mm width) with sterile razorblades and pooled to 100mg per sample. DNA was extracted from the pooled midribs per plant using PowerPlant^TM^ DNA Isolation Kit (MoBio, CA, USA) following instructions of the manufacturer.

### Real time PCR

Taqman real-time PCR using primers HLBas and HLBr and probe HLBp were performed on all plant samples as described by Li et al. (2006). An additional primer-probe set and probe based on plant cytochrome oxidase (COX) were used as positive internal control (Li et al. 2006). The real-time PCR amplifications were performed in a LightCycler^®^ 480 Real-time PCR Instrument (Roche Diagnostics, IND, USA). All real-time PCR reactions were run for one cycle at 50°C for 2 min, one cycle at 95°C for 20 s, 35 cycles at 95°C for 1 s and 58°C for 40 s. All reactions were performed in triplicate and each run contained three Las-positive and three Las-negative citrus plants as controls (obtained from a BSL3 greenhouse at the DPI, Gainesville, Florida, USA). The data were analyzed using the LightCycler^®^ 480 Software release 1.5.0. The number of cycles was limited to 35 to avoid occurrences of false-positive signals (Sipos et al. 2007).

### *Candidatus* Liberibacter spp. 16S rDNA gene PCR

Amplicons for *Ca.* Liberibacter spp.16S rDNA gene were generated using the PCR method adapted from Jagoueix et al. (1994) and Jagoueix et al. (1996). Triplicate PCR reactions were performed per plant extract per annealing temperature, 1 µL was added to 49 µL PCR reaction mixtures consisting of Tris–HCl (pH 8.3),10 mM; KCl, 10 mM; MgCl2, 2.5 mM; each deoxyribonucleoside triphosphate, 200 mM; 400 mM of each primer, OI1 (5’-GCGCGTATGCAATACGAGCGGCA-3’) and OI2c (5’-GCCTCGCGACTTCGCAACCCAT-3’), and 5 U of *Taq* DNA Polymerase (New England BioLabs Inc., USA). PCR amplifications were run in an Applied Biosystems Veriti^TM^ Thermal Cycler (Applied Biosystems, USA) programmed at one cycle of 95°C, 5 min; 35 cycles at 95°C for 40 s, 54°C for 30 s, and 72°C for 90 s; and one cycle 72°C for10 min. The primer set OI1 and OI2c was designed to amplify sequences of Las. To use the same primer set to amplify sequences of a wider range of *Ca.* Liberibacter species we performed the PCR as described above while diminishing the annealing temperature 0.5°C down to 48°C. To search for amplicons with expected size (approximately 1160 bp), all pseudo triplicates of the 130 plant DNA extract PCR reactions were analyzed in 1.5% agarose gels, run at 75V per 1.5h and stained with ethidium bromide for each of the annealing temperatures analyzed. The PCR products obtained were used for cloning.

### *Candidatus* Liberibacter spp. rplA/rplJ gene PCR

Amplicons for *Ca.* Liberibacter spp. rplA/rplJ gene were generated using the PCR method adapted from Hocquellet et al. (1999). Triplicate PCR reactions were performed for plant extracts per annealing temperature, 1 µL was added to 49 µL PCRreaction mixtures consisting of Tris–HCl (pH 8.3),10 mM; KCl, 10 mM; MgCl2, 2.5 mM; each deoxyribonucleoside triphosphate, 200 mM; 400 mM of each primer, A2 (5’-TATAAAGGTTGACCTTTCGAGTTT-3’) and J5 (5’-ACAAAAGCAGAAATAGCACGAACAA-3’), and 5 U of *Taq* DNA Polymerase (New England BioLabs Inc., USA). PCR amplifications were run in an Applied Biosystems Veriti^TM^ Thermal Cycler (Applied Biosystems, USA) programmed at one cycle of 95°C, 5 min; 35 cycles at 92°C for 20 s, 62°C for 20 s, and 72°C for 45 s; and one cycle 72°C for 5 min. The primer set A2 and J5 was designed to amplify sequences of Las. To use the same primer set to amplify sequences of *Ca.* Liberibacter species we performed the PCR as described above and diminishing the annealing temperature 0.5°C in different PCR runs, down to 56°C. To search for amplicons with the expected size (approximately 1160 bp), all pseudo triplicates of the 130 plant DNA extract PCR reactions were analyzed in 1.5% agarose gels, run at 75V per 1.5h and stained with ethidium bromide for each of the annealing temperatures analyzed. The PCR products obtained were used for cloning.

### Construction of a 16S rRNA and rplA/rplJ gene clone library made from *Candidatus* Liberibacter spp

A 16S rRNA gene and a rplA/rplJ clone library was made from midrib-DNA extracts of each plant that had tested positive in real time PCR with the primers HLBas and HLBr. Hence, purified DNA extracts were PCR amplified using primer sets OI1 / OI2c and A2 / J5. The PCR products with the expected fragment size were purified from non-incorporated dNTPs and primers using the QIAquick PCR purification kit (Qiagen, Hilden, USA). Purified PCR product was cloned into the pCR2.1-TOPO vector from the TOPO-TA PCR cloning kit (Invitrogen, USA) and introduced into *Escherichia coli* Top10cells (Invitrogen, USA) by transformation according to the protocol provided by the manufacturer. White colonies, indicating insertional inactivation of the lacZ gene, were PCR amplified with primers annealing with the M13F andM13R sites in the pCR2.1-TOPO vector using the procedure provided by the manufacturer. In total, seven clones for each primer set for each plant with the expected fragment size were selected for later sequencing.

### Sequencing of PCR fragments

DNA sequencing of the samples was done at the University of Florida DNA Sequencing core Laboratory. Sequencing reactions were performed using ABI Prism BigDye Terminator cycle sequencing protocols (part number 4337036) developed by Applied Biosystems (Perkin-Elmer Corp., Foster City, CA). ABI prism BigDye Terminators v.1.1 cycle sequencing reactions were assembled in 20µl reaction volume by adding 25ng DNA, 10 pmols primer, 2µl of BigDyer terminator, 3µl of 5X sequencing buffer and 5% DMSO. The cyclic profile was performed as recommended by the manufacturer. The excess dye-labeled terminators were removed using MultoScreen® 96-well filtration system (Millipore, Bedford, MA, USA). The purified extension products were dried in SpeedVac^®^ (ThermoSavant, Holbrook, NY, USA) and then suspended in Hi-di formamide. Sequencing reactions were performed using POP-7 sieving matrix on 50-cm capillaries in an ABI Prism^®^ 3130 Genetic Analyzer (Applied Biosystems, Foster City, CA, USA) and were analyzed by ABI Sequencing Analysis software v. 5.2 and KB Basecaller.

### Phylogenetic Analysis

Sequence data from all clones were first checked for chimeras using the Bellerophon (version 3) (Huber et al. 2004), and non-chimeric sequences were used for phylogenetic analysis. Phylogenetic trees were constructed using the gene sequences of *Ca*. Liberibacter spp. and *Liberibacter crescens* in Genbank (Table S2). Phylogenetic distances of the partial 16S rDNA gene sequences were calculated using the ARB software package (Kumar et al. 2006; Ludwig et al. 2004) optimized with X model (e.g. uncorrected P distance or may be HKY85) using the SILVA database Ref 106 with a 99% criterion applied to remove redundant sequences (Pruesse et al. 2007) from the seven plants positive for the PCR with primer set OI1 and OI2c. Aligned sequences were manually edited taking 16S rRNA sequence secondary structures into consideration. Reconstruction of phylogenetic relationships was based on neighbor joining (Ludwig et al.1998). The branches were tested with bootstrap analysis (1,000 iterations). The seven isolates were so similar to each other that they were considered as one strain: *Ca*. Liberibacter sp. strain UFEPI.

**Table 2.** 16S rDNA gene pairwise similarity among *Candidatus* Liberibacter spp.

| # | *Candidatus*<br>Liberibacter | Accession<br>number | 1<sup>a</sup> | 2 | 3 | 4 | 5 | 6 | 7 | 8 | 9 | 10 | 11 | 12 | 13 | 14 | 15 | 16 | 17 | 18 |
| --- | --- | --- | --- | --- | --- | --- | --- | --- | --- | --- | --- | --- | --- | --- | --- | --- | --- | --- | --- | --- |
| 1 | terrae str.<br>UFEPI | MF463003 |  |  |  |  |  |  |  |  |  |  |  |  |  |  |  |  |  |  |
| 2 | unnamed sp.<br>M-20 | GU553033 | **0.025** |  |  |  |  |  |  |  |  |  |  |  |  |  |  |  |  |  |
| 3 | asiaticus | CP001677 | **0.030** | 0.004 |  |  |  |  |  |  |  |  |  |  |  |  |  |  |  |  |
| 4 |  | EU921615 | **0.030** | 0.004 | 0.002 |  |  |  |  |  |  |  |  |  |  |  |  |  |  |  |
| 5 |  | EU644449 | **0.032** | 0.006 | 0.004 | 0.0 |  |  |  |  |  |  |  |  |  |  |  |  |  |  |
| 6 |  | EU921617 | **0.029** | 0.005 | 0.003 | 0.003 | 0.005 |  |  |  |  |  |  |  |  |  |  |  |  |  |
| 7 |  | EU921616 | **0.032** | 0.006 | 0.004 | 0.004 | 0.006 | 0.005 |  |  |  |  |  |  |  |  |  |  |  |  |
| 8 |  | FJ750456 | **0.032** | 0.006 | 0.004 | 0.004 | 0.006 | 0.005 | 0.006 |  |  |  |  |  |  |  |  |  |  |  |
| 9 |  | FJ750459 | **0.034** | 0.008 | 0.006 | 0.006 | 0.008 | 0.007 | 0.008 | 0.008 |  |  |  |  |  |  |  |  |  |  |
| 10 |  | L22532 | **0.027** | 0.003 | 0.001 | 0.001 | 0.003 | 0.001 | 0.003 | 0.003 | 0.005 |  |  |  |  |  |  |  |  |  |
| 11 | africanus | EU921619 | **0.046** | 0.024 | 0.021 | 0.021 | 0.024 | 0.023 | 0.024 | 0.024 | 0.024 | 0.020 |  |  |  |  |  |  |  |  |
| 12 |  | EU754741 | **0.050** | 0.028 | 0.026 | 0.026 | 0.028 | 0.027 | 0.028 | 0.028 | 0.028 | 0.026 | 0.013 |  |  |  |  |  |  |  |
| 13 |  | L22533 | **0.041** | 0.019 | 0.017 | 0.017 | 0.020 | 0.017 | 0.019 | 0.019 | 0.019 | 0.016 | 0.004 | 0.009 |  |  |  |  |  |  |
| 14 | psyllaorous | EU812557 | **0.044** | 0.026 | 0.024 | 0.022 | 0.026 | 0.025 | 0.026 | 0.026 | 0.028 | 0.022 | 0.027 | 0.028 | 0.022 |  |  |  |  |  |
| 15 | solanacearum | EU935004 | **0.044** | 0.026 | 0.024 | 0.021 | 0.026 | 0.025 | 0.026 | 0.026 | 0.028 | 0.022 | 0.027 | 0.028 | 0.022 | 0.000 |  |  |  |  |
| 16 | americanus | EU921624 | **0.058** | 0.043 | 0.041 | 0.038 | 0.043 | 0.042 | 0.043 | 0.043 | 0.043 | 0.038 | 0.039 | 0.044 | 0.034 | 0.041 | 0.041 |  |  |  |
| 17 |  | AY742824 | **0.056** | 0.042 | 0.039 | 0.037 | 0.042 | 0.041 | 0.042 | 0.042 | 0.042 | 0.037 | 0.038 | 0.043 | 0.033 | 0.039 | 0.039 | 0.001 |  |  |
| 18 |  | FJ263691 | **0.057** | 0.043 | 0.040 | 0.038 | 0.043 | 0.042 | 0.043 | 0.043 | 0.043 | 0.038 | 0.039 | 0.044 | 0.034 | 0.041 | 0.041 | 0.002 | 0.001 |  |
| 19 | europaeus | FN678792 | **0.049** | 0.036 | 0.033 | 0.034 | 0.036 | 0.035 | 0.036 | 0.036 | 0.038 | 0.033 | 0.036 | 0.040 | 0.032 | 0.042 | 0.042 | 0.032 | 0.031 | 0.032 |

<sup>a</sup> in bold pairwise similarity of *Candidatus* Liberibacter terrae and other *Ca.* Liberibacter spp.

Closest matches to the sequences of the partial rplA/rplJ gene were obtained by BlastN search. Sequences were aligned. Phylogenetic distances of partial rplA/rplJ gene for the same plants positive for the OI1 and OI2c primer set and their closest matches were calculated using the software Mega 5 optimized with X model (Tamura et al. 2011). Reconstruction of phylogenetic relationships was based on neighbor joining (Ludwig et al.1998). The branches were tested with bootstrap analysis (1,000 iterations).

### Nucleotide sequence accession numbers

The DNA sequences of the partial 16S rRNA genes (1,049 bp) of seven *Ca.* Liberibacter sp. strains UFEPI and of the rplA/rplJ gene (739 bp) of one strain (1R1T9) of *Ca.* Liberibacter sp. strain UFEPI were deposited in the EMBL Nucleotide Sequence Database (Cochrane et al. 2009) under accession numbers MF 463003 to MF463009, and MK125061, respectively.

## RESULTS

### Disease symptoms and plant growth

Four months after planting, eight seedlings in non-autoclaved soil from grove B showed pronounced chlorosis, while their counterparts in autoclaved soil did not show HLB-like symptoms (Fig. S1). Severe yellowing and stunting symptoms that can be associated with HLB (Folimonova et al. 2009) were observed on two plants in non-autoclaved soil from grove B 8.5 months after planting (Fig. 1 D and F). These symptoms were not observed on plants in soil from grove A, nor on those in autoclaved soil from grove B or in pasteurized potting mix.

After 2.5 months, plant heights were significantly (P<0.05) greater in autoclaved than in nonautocalved soils according to paired t-tests, while there were no differences in plant height between soil from grove A versus grove B (Fig. 1B and Table 1). The final plant weights after 8.5 months were also significantly (P<0.05) higher in autoclaved than in non-autoclaved soil, without grove differences (Table 1).

### Real-time PCR with primers HLBas / HLBr and probe HLBp

Las group-specific Real time PCR was performed as previously described using HLBas and HLBr primers as well as COXf and COXr as internal controls (Li et al. 2006). The Ct values of the negative control samples (from disease-free plants maintained at the DPI) were always >40. The Ct values of the positive controls (HLB symptomatic plants in a BSL3 greenhouse at the DPI) varied from 15.59 to 25.12. Seven mandarin plants grown in non-autoclaved soil B (1AR1T9, 1AR4T1, 2AR2T2, 4AR4T1, 5AR1T9, 2BR2T2 and 4BR4T1) showed Ct values that ranged from 29.5 to 33.8 (Table S3). The first number stands for the plant number in the greenhouse, the capital letters A or B for the time of sampling in the greenhouse, and the capital letters R and T stand for row and tree numbers in the grove. Thus, five of the positive plant extracts (1AR1T9, 1AR4T1, 2AR2T2, 4AR4T1 and 5AR1T9) were recovered from plants that were sampled 2.5 months after the beginning of the greenhouse experiment and two of them (2BR2T2 and 4BR4T1) were sampled after 8.5 months. The soil where these plants were grown came from three different locations (R1T9, R2T2, and R4T1) inside grove B.

**Table 3.** β-operon pairwise similarity among *Candidatus* Liberibacter spp.

| # | *Candidatus* | Accession | 1<sup>a</sup> | 2 | 3 | 4 | 5 | 6 |
| --- | --- | --- | --- | --- | --- | --- | --- | --- |
|  | Liberibacter | number |  |  |  |  |  |  |
| 1 | terrae str. UFEPI |  |  |  |  |  |  |  |
| 2 | asiaticus | GU074017 | **0.124** |  |  |  |  |  |
| 3 | asiaticus | GQ890156 | **0.124** | 0.000 |  |  |  |  |
| 4 | asiaticus | FJ394022 | **0.124** | 0.000 | 0.000 |  |  |  |
| 5 | africanus | U09675 | **0.218** | 0.237 | 0.237 | 0.237 |  |  |
| 6 | africanus | GU120043 | **0.218** | 0.237 | 0.237 | 0.237 | 0.000 |  |
| 7 | americanus | EF122254 | **0.478** | 0.469 | 0.469 | 0.469 | 0.461 | 0.461 |

<sup>a</sup> in bold pairwise similarity of *Candidatus* Liberibacter terrae str. UFEPI and other *Ca.* Liberibacter spp.

### *Candidatus* Liberibacter spp. 16S rDNA gene analysis

None of the DNA extracts separately extracted from the 130 plants in the greenhouse experiment produced amplicons with the expected fragment size (approximately 1160 bp) with the primers OI1 and OI2c using 62°C as annealing temperature. After this initial test we performed different runs of PCR with the same samples and primers but for each round we diminished the annealing temperature by 0.5°C. Seven samples (1AR1T9, 1AR4T1, 2AR2T2, 4AR4T1, 5AR1T9, 2BR2T2 and 4BR4T1) generated single band amplicons with approximately the expected fragment size when the PCR annealing temperature was 59.5°C, below the recommended temperature for Las (Fig. S2). The samples that showed a single band of the approximate expected size in an agarose gel were the same that were positive for the real time PCR with primers HLBas / HLBr and probe HLBp.

After cloning and sequencing of each individual band and assemblage of the different contigs the fragment size of the positive samples were approximately 1,049 bp. These sequences did not have the expected fragment size for Las (1,160 bp). This data indicated that the bacteria detected with primers OI1 and OI2c using 52.5°C as annealing temperature were not Las. Analysis of these 16S rDNA gene sequences with the program Bellerophon (version 3) indicated that all of them were non-chimeric. The software package ARB and the SILVA database Ref 106 with a 99% criterion applied to remove redundant sequences were used to further analyze the phylogeny of the partial 16S rDNA sequences recovered in this study. Matrix analysis using this software package revealed that the seven 16S rDNA gene sequences obtained in this study were more than 99.8% similar and therefore considered from the same species (Table S4). Moreover, all seven partial 16S rDNA sequences were most closely affiliated with *Ca.* Liberibacter spp. (Table 2) and initially we named these sequences *Ca.* Liberibacter sp. str. UFEPI. The closest relative to the partial 16S rDNA sequences found in this study was the uncultured bacterium clone M-20 (96.6% similarity) recovered from endosymbiontic bacteria in a *Diaphorina citri* (Tian et al. 2010 – direct submission to Genbank, accession number GU553033). *Ca.* Liberibacter sp. str. UFEPI was on average 95.9% similar to Las, 94.8% similar to Laf, 94.5% similar to *Ca.* Liberibacter psyllaurous and *Ca.* Liberibacter solanacearum and 93.6% similar to Lam (Table 2). A phylogenetic tree without outgroup shows that the uncultured clone M-20 is indeed most closely related to *Ca*. Liberibacter sp. str. UFEPI (Fig. 2A and Fig. S3). However, when *Bradyrhizobium japonicum* is used as outgroup, clone M-20 is closely related to Las, but not to *Ca*. Liberibacter sp. str. UFEPI (Fig. 2B). Similarly, when *Liberibacter crescens* is included, clone M-20 is related more closely to Las than to str. UFEPI (Fig. S4).

**Fig. 2.**
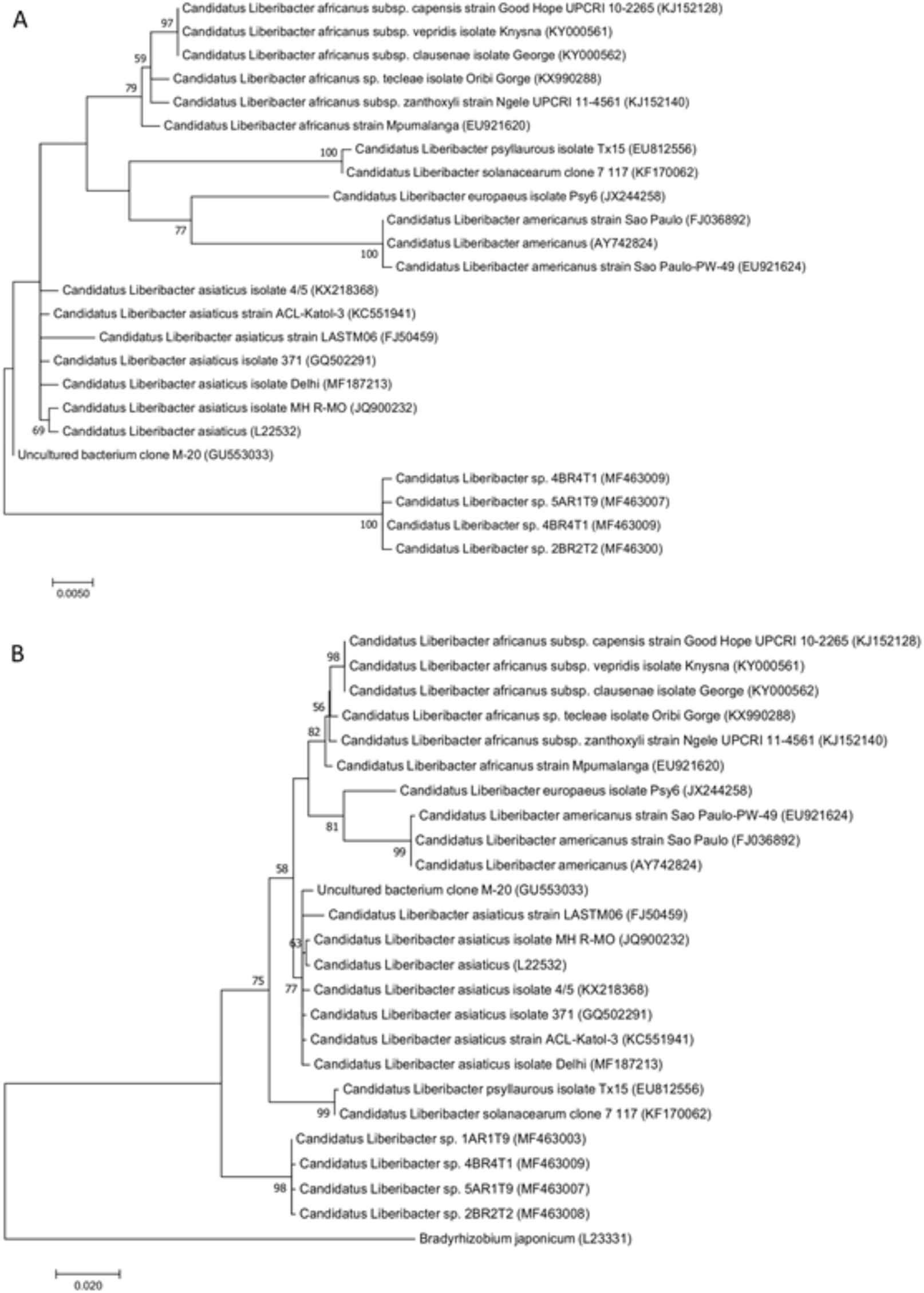
Phylogenetic trees based on maximum-likelihood analysis of the 16S rDNA sequences with various members within the genus *Candidatus* Liberibacter without outgroup (**A**) and with *Bradyrhizobium japonicum* as outgroup (**B**). Phylogeny was inferred using the Tamura-Nei model (Tamura and Nei 1993) with gamma correction to account for site variations. Bootstrap values based on 1,000 replications are shown at branch nodes and values under 50 are not shown. Genbank accessions are shown on the tree for sequences indicated in brackets. Bar, 0.020 substitutions per nucleotide position.

### *Candidatus* Liberibacter spp. β operon (gene) analysis

Similar to what was observed for the Las 16S rDNA gene group-specific primers, none of the 130 plant DNA extracts generated a single band with the expected fragment size (703 bp) with primers A2 and J5. These data confirm that the *Ca.* Liberibacter sp. str. UFEPI may not be Las. To obtain a single band with approximately the expected amplicon size produced by the primers A2 and J5 we employed the same strategy used for the 16S rDNA gene. After diminishing the annealing temperature of PCR reactions with primers A2 and J5 to 59.5°C, single band amplicons with approximately the expected fragment size were observed in agarose gels in the same samples that were positive for real time PCR and with primers OI1 and OI2c (1AR1T9, 1AR4T1, 2AR2T2, 4AR4T1, 5AR1T9, 2BR2T2 and 4BR4T1).

Single clones of each of the seven different DNA extracts positive for β operon products at 59.5°C were cloned and sequenced. After assembling the different contigs it was demonstrated that the amplicons obtained in this study were 739 bp in length. No chimeras were detected among these sequences. These differ from the A2 and J5 amplicon fragment expected for Las (703 bp) and Lam (669 bp). The software Mega5 (Tamura et al. 2011) was used to determine the pairwise distance among the seven β operon products obtained in this study and among their closest relatives in the NCBI database. It was demonstrated that the different β operon sequences from this study were more than 99.6% similar and therefore they are considered from the same bacterium species (Table S5). Phylogenetic analysis of the *Ca.* Liberibacter sp. str. UFEPI β operon (Fig. 3) demonstrated that the closest relatives of this bacterium were Las strains (accession numbers FJ394022, GQ890156 and GU074017) approximately 87.6% similarity, followed by Laf strains (U09675 and GU120043) approximately 78.2% similarity (Table 3). The β operon gene of the uncultured bacterium clone M-20 was not available in Genbank. Phylogenetic analysis of the multilocus sequences of *Ca*. Liberibacter species based on maximum likelihood of two concatenated loci, 16S rRNA and rplA/J (a total of 1788 bp), demonstrated again that Ca. Liberibacter strain UFEPI formed a separate branch most closely related to Las (Fig. 4).

**Fig. 3.**
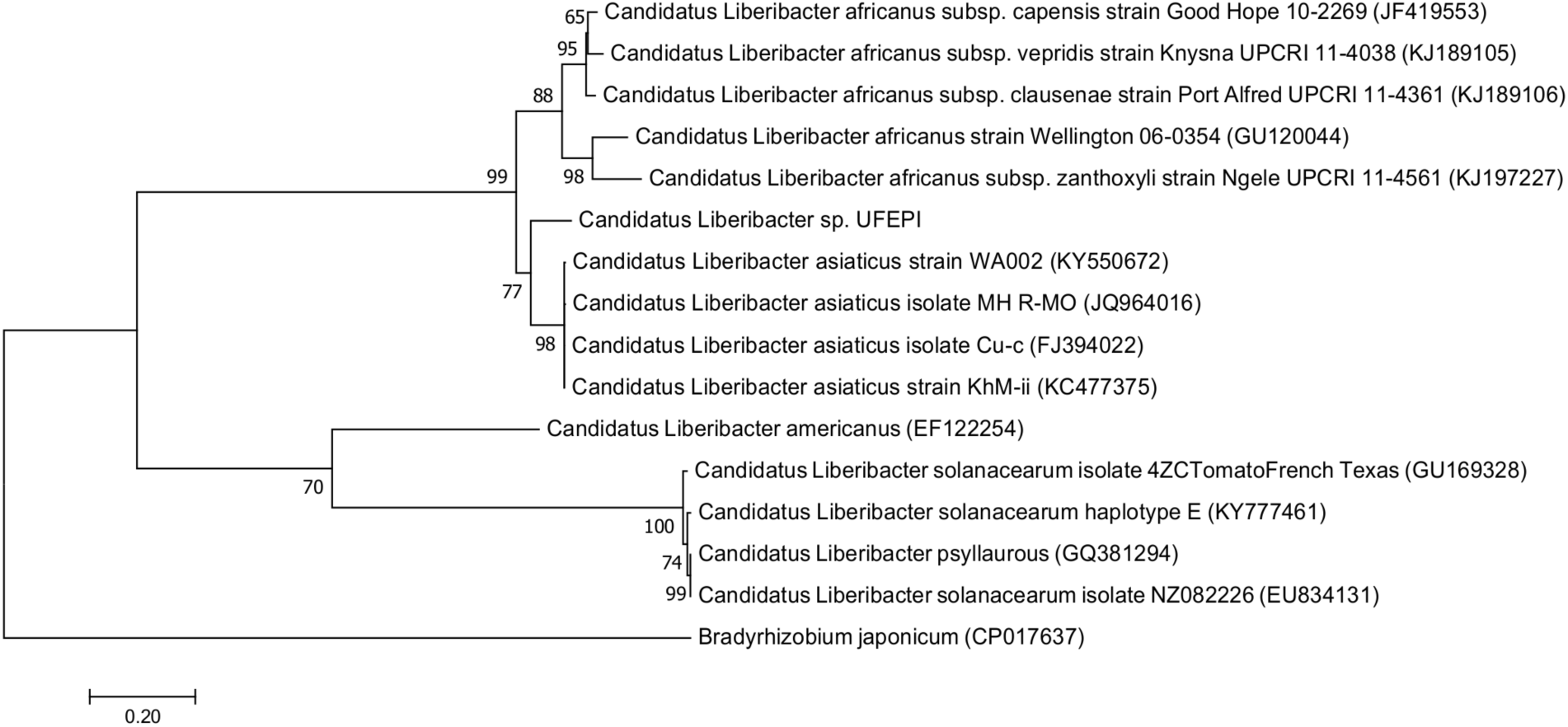
Phylogeny based on maximum-likelihood analysis of the rplA/J gene sequences (of strain UFEPI-R1T9) with various members within the genus *Candidatus* Liberibacter and *Bradyrhizobium japonicum* as outgroup. Phylogeny was inferred using the Tamura-Nei model (Tamura and Nei 1993) with gamma correction to account for site variations. Bootstrap values based on 1,000 replications are shown at branch nodes. Genbank accessions are shown on the tree for sequences indicated in brackets. Bar, 0.20 substitutions per nucleotide position.

**Fig. 4.**
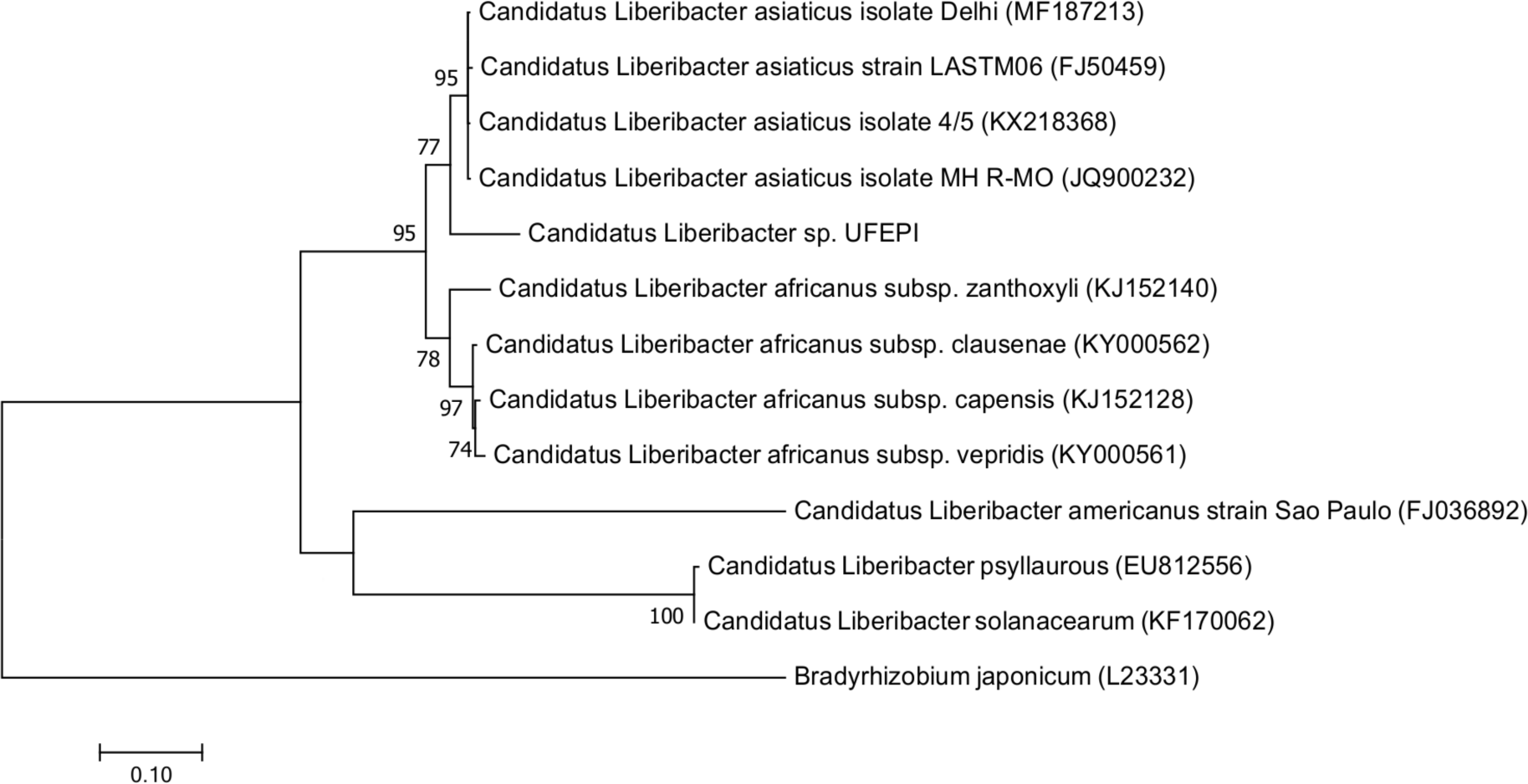
Phylogenetic analysis of the multilocus sequences of Candidatus Liberibacter species based on maximum-likelihood analysis of two concatenated loci, 16S rRNA and rplA/J (a total of 1788 bp). *Bradyrhizobium japonicum* was used as outgroup. Phylogenetic tree was generated in MEGA7 using the mura-Nei model (Tamura and Nei 1993) with gamma correction to account for site variations. Bootstrap values based on 1,000 replications are shown at branch nodes and values under 50 are not shown. Genbank accessions are shown on the tree for sequences indicated in brackets. Bar, 0.10 substitutions per nucleotide position.

## DISCUSSION

Our research provides the first evidence that a new *Ca.* Liberibacter strain (UFEPI) different from those currently known to be associated with HLB (i.e. Las, Laf and Lam) can reside in citrus trees displaying HLB-like symptoms (Folimonova et al. 2009), including yellowing and stunting of plants but no leaf mottling. Our data also indicate that this novel *Ca.* Liberibacter strain is not necessarily transmitted to citrus trees by psyllids as our experiments were carried out in an insect-free greenhouse in an area that was still free from ACP and HLB at the time of the experiments (Shen et al. 2013b). Yet, the closest relative of strain UFEPI was an unnamed, uncultured bacterium M-20 obtained from *D. citri* (Tian et al. 2010 – direct submission to Genbank, accession number GU553033). Another relevant finding in this study was that all the plant samples where the new *Ca.* Liberibacter strain UFEPI was detected originated from the same treatment, non-autoclaved soil B. It was not detected in plants in autoclaved soils or pasteurized potting mixture. Moreover, the same *Ca.* Liberibacter strain occurred in samples collected under three of the five different citrus trees in one HLB-positive grove, indicating that the transmission of the new *Ca.* Liberibacter strain is not a rare event. Considering these observations, the question arises what the mode of transmission of the new *Ca.* Liberibacter strain was to the mandarin plants. As all the plants were grown in an insect free greenhouse facility we have to exclude transmission by psyllids in this experiment. That leaves three alternatives: (i) the new *Ca.* Liberibacter strain is an endophyte or pathogen that is seed transmitted; (ii) this bacterium can be transmitted via the soil plant interface or (iii) it can be transmitted by an unidentified soilborne vector.

If seed transmission would be the case, most likely we would have detected the new *Ca.* Liberibacter strain UFEPI in the mandarin trees that were planted in soil A, in autoclaved soil B and in potting mixture. Still considering seed transmission, one could hypothesize that *Ca.* Liberibacter strain UFEPI was transmitted by seed and required the biotic and abiotic conditions found in soil B to be expressed in the plants. Several studies have demonstrated that Las, the closest recognized relative of *Ca.* Liberibacter strain UFEPI, can be detected in seeds but movement into the growing plant is so rare that Las is considered not to be seed-borne (Albrecht and Bowman 2009; Hartung et al. 2010; Hilf 2011). Moreover, the mandarin seedlings used for our experiment originated from a reputable, commercial, enclosed citrus nursery, and were unlikely to be infected by Las or a closely related strain.

Therefore, the most likely source of this *Ca.* Liberibacter strain was the soil collected from HLB-infested citrus grove B, as it was not found in citrus seedlings grown in soil from grove A, in either of the autoclaved soils or in pasteurized potting mix. The potential soil-borne nature of HLB was thought to be non-credible when preliminary data were first presented at the Second International Research Conference on Huanglongbing in 2011 (Nunes da Rocha et al. 2011). One of the arguments for this skepticism was that Las could not possibly survive in soil due to its very small genome, missing the genes for several essential enzymes including those for glycolysis (Duan et al. 2009; Hartung et al. 2011; Jain et al. 2017). In the meantime however, several other observations suggest the potential soil-borne nature of Las and related *Ca*. Liberibacter spp. Las can move both basipetally and acropetally in the phloem depending on the sink of the carbon compounds, but the presence of high densities of Las in roots before the pathogen is found in the foliage of citrus trees (Johnson et al. 2014) suggests that root uptake could possibly take place besides transmission by psyllids. Moreover, phloem decline occurs in young roots before older roots (Kumar et al. 2018). Another observation supporting the potential soil-borne nature of HLB is the high frequency of reinfection of newly planted disease-free young citrus trees at the locations where infected trees were removed, despite intensive vector control in those groves (Timmer 2014). Anaerobic soil disinfestation after removal of HLB-symptomatic trees delayed the appearance of HLB symptoms in young transplants (Rosskopf, personal communication). These observations remind us of the well-documented replant diseases in other fruit trees (Browne et al. 2018; Mazzola and Manici 2012; Yang et al. 2012). These diseases can be caused by a variety of pathogens, depending on the pathogen that was present in the roots of the removed trees. Recently, we detected Las in deep-sequenced 16S rDNA from colony mixtures isolated from citrus rhizosphere soil on a low-carbon agar medium (Ascunce et al, in preparation). Finally, *Ca*. Liberibacter solanacearum haplotype C was detected in stolons and tubers of field-grown asymptomatic potato plants in infected carrot fields (Haapalainen et al. 2018a). The below-ground potato materials were not in contact with psyllids, and carrot psyllids could not transmit haplotype C to potato leaves under controlled conditions (Haapalainen et al. 2018a). Thus, also in this case, some form of soil transmission could have been possible.

Although these arguments support the potential soil-borne nature of HLB, we are uncertain how the new strain of *Ca*. Liberibacter sp. could have crossed from the soil environment into the plants. There are several possible pathways. Young roots grow preferentially towards and into decaying roots, which may still carry the cells of a pathogen (Caldwell et al. 1991; McKee 2002; van Vuuren et al. 1996). The pathogen may then enter the young roots through natural openings, wounds or the root tip. In our experiment, the soil was sieved but cut root pieces of HLB-positive trees were returned to the soil, and the roots of the seedlings could have picked up *Ca*. Liberibacter str. UFEPI, even if it could not multiply in soil due to a lack of essential enzymes (Jain et al. 2017). Alternatively, the pathogen could have been vectored by contaminated soilborne insect vectors, nematodes or fungi, which could have transferred the pathogen to young roots.

Insects that spend (part of) their lifecycle on roots and in soil could transmit plant pathogens from one host crop to the next in a particular area. Mealy bugs can be associated with soil during the first larval stage, the crawler stage, when they move to a new feeding site (Naidu et al. 2014; Osborne, 2016). Some mealy bug species are specifically adapted to feeding on roots, the so-called root mealy bugs (*Geococcus coffeae*, *Rhizoecus* spp.), which occur in Florida next to various other mealybug species, such as the citrus mealybug (Dekle, 1965; Osborne, 2016). *G. coffeae* has been found on Chinese Boxorange (Dekle, 1965), a close relative of citrus that can harbor Las (Hung et al. 2001). Several other mealybug species of the genus *Pseudococcus* can survive on roots in the soil and transmit, for example, Grapevine Leafroll Virus from remnant roots in uprooted vineyards to new plantings (Bell et al. 2009; Naidu et al. 2014). At least one (foliar) mealy bug species, *Ferrisia virgata*, was found carrying Las, although it did not transmit HLB when tested (Pitino et al. 2014). Thus, *Ca*. Liberibacter strain UFEPI could possibly have been transmitted from soil to the young mandarin plants by (root) mealybugs that were not observed during our experiment. In addition, root knot nematodes could have been involved in transmission of *Ca*. Liberibacter cells into citrus roots, as these nematodes are intimately associated with the phloem of infected plants (Absmanner et al. 2013; Bartlem et al. 2014). Further studies are necessary to address questions about the potential modes of soil transmission by various *Ca*. Liberibacter species.

The novel *Ca.* Liberibacter sp. found in this study was initially detected using the same primers used for real time quantification of Las developed by Li et al. (2006). This protocol and its optimized version (Li et al. 2008) are the most used primer-sets for the confirmation of HLB. The real-time PCR CT values obtained for our citrus leaf DNA extracts ranged from 29.5 to 33.8, indicating that we had detected Ca. Liberibacter sp.. These CT values were not sufficient to indicate that our citrus trees were contaminated with Las (Shin and van Bruggen 2018). In another experiment, CT values between 31 and 36 were obtained using the same primer set for citrus samples that contained *Bradyrhizobium*, which is closely related to *Ca*. Liberibacter (Shin and van Bruggen 2017). For samples with high CT values, and therefore low titers of the target bacterium, it is suggested that a second detection method be used to determine if plant samples are infected with *Ca.* Liberibacter spp. associated with citrus HLB (Tatineni et al. 2008).

Las group-specific 16S rDNA gene detection, according to Jagoueix et al. (1994) and Jagoueix et al. (1996), demonstrated that the bacterium detected in our samples by real-time PCR was closely related to Las, but different enough to warrant further identification. The CT values observed in the real-time PCR indicated that the titer of the bacterium that produced the signal was low. Instead of sequencing a large number of clones made with general bacterial primers, we diminished the stringency of the annealing temperature of the primers OI1 and OI2c. A single amplicon with size similar to that expected for Las was detected using 52.5°C as annealing temperature. In this amplicon sites are present for annealing with the primers HLBas and HLBr and the probe HLBp (data not shown). This may explain why this bacterium was detected by the Las group-specific real-time PCR. Sequencing and phylogenetic analysis demonstrated that *Ca.* Liberibacter sp. str. UFEPI is a novel *Ca.* Liberibacter species closely related to Las. To increase the confidence of these findings and to generate more genetic information about *Ca.* Liberibacter sp. str. UFEPI we sequenced the β operon of this bacterium using the primers A2 and J5 previously designed to amplify the β operon fragment of Las (Hocquellet et al. 1999). No amplicon was detected using 62°C as annealing temperature but single band amplicons with approximately the expected size were detected when using 59.5°C as annealing temperature. Phylogenetic analysis demonstrated that these sequences were also shared with the *Ca.* Liberibacter group.

The combined findings of this study demonstrate that *Ca.* Liberibacter sp. str. UFEPI belongs to the HLB-associated *Ca*. Liberibacter group and is likely transmitted to citrus plants through the soil. Other novel strains belonging to the *Ca.* Liberibacter group have been described recently (Gupta et al. 2012; Keremane et al. 2015; Morris et al. 2017; Roberts et al. 2015; Tian et al. 2010). *Ca.* Liberibacter sp. str. UFEPI is 96.6% similar to an unnamed *Ca.* Liberibacter sp. M-20 (accession number GU553033) isolated from a psyllid (*D. citri*) in China (Tian et al. 2010 – direct submission to Genbank), 95.9% similar to Las, 94.8% similar to Laf, 94.5% similar to *Ca.* Liberibacter psyllaurous and *Ca.* Liberibacter solanacearum and 93.6% similar to Lam. These results indicate that *Ca*. Liberibacter str. UFEPI represents a new species for which the name *Ca.* Liberibacter terrae is putatively proposed. Naming this novel species will prevent misclassification of species that are similar to this bacterium but differ from the other recognized species of *Ca*. Liberibacter. *Ca.* Liberibacter terrae likely can enter citrus plants from soil and induce part of the syndrome typical for HLB. It is not known if *Ca.* Liberibacter terrae can be transmitted by psyllids or other insect vectors and if it would result in yield loss. Previously, different strains of Las were shown to induce some of the characteristic HLB symptoms but not all (Tsai et al. 2008). Thus, HLB may be a disease complex caused by a number of closely related *Ca*. Liberibacter species.

Our findings could have important practical implications. The realization that the commonly accepted ecological cycle of *Ca*. Liberibacter spp. between psyllids and plant phloem may be too narrow and may include a soil phase (for example: from psyllids to leaves and in infected aborted leaves to soil, plant roots and phloem; or from psyllids via honeydew secretions to soil, plant roots and phloem; or from phloem in plant roots to nematodes or mealybugs and back to phloem) could have major consequences for disease management. Current methods to combat HLB primarily rely on insect vector control, cutting of diseased trees and replanting with pathogen-free new trees (Bové 2014). Models to predict the spread of HLB and effectiveness of various control measures have not considered the possibility of reinfection of young trees from the roots of HLB-infected trees that were cut to reduce the inoculum availability (Chiyaka et al. 2012; Craig et al. 2018; Lee et al. 2015; Luo et al. 2017; Taylor et al. 2016), and may need to be adjusted.

### Conclusions and future perspective

We detected a novel species in the Liberibacter group, putatively called *Ca.* Liberibacter terrae. This bacterium was detected in plants grown in non-autoclaved soil B and was not detected in those plants that grew in non-autoclaved soil A, autoclaved soil A or B and pasteurized potting mixture. Although this bacterium belongs to the *Ca*. Liberibacter group it was likely transmitted to the citrus plants through the soil-plant interface. This trait is very different from the other recognized species of this group that are transmitted to plants by psyllids. Additional studies will need to be undertaken that will determine how this bacterium crosses the soil-plant interface. Moreover, it will be important to determine if *Ca.* Liberibacter terrae is a phloem-residing bacterium that could also be transmitted by psyllids. Our data indicate that more ecological studies are necessary to find the actual diversity of species that belong to the *Ca.* Liberibacter group. A better understanding of the ecology and epidemiology of this new species may shed light on improvements in the management of plant diseases associated with various *Ca*. Liberibacter species, including huanglongbing. Finally, we suggest additional research to investigate if the well-established *Ca*. Liberibacter species associated with HLB (Las, Laf and Lam) could possibly be soil-borne.

## ACKNOWLEDGEMENTS

Partial financial contributions came from the Smallwood Foundation, the Esther B. O’Keeffe Foundation, the Emerging Pathogens Institute of the University of Florida (UFEPI), and the Institute of Food and Agricultural Sciences (IFAS) of the University of Florida. The authors would like to thank Ellen Dickstein for assistance setting up the experiments, the grove managers who supplied the soil for this study, and Debra Jones of the Department of Plant Industry (Gainesville, FL) for providing citrus leaves positive and negative for Las that were used as controls in all PCR reactions performed in this study.

## COMPLIANCE WITH ETHICAL STANDARDS

***Conflict of Interest.*** The authors declare that they have no conflict of interest.

## Supplement

### Materials and Methods

#### Soil samples

Soil was collected from two citrus groves in central Florida: (A) the USDA Citrus Research Station in Winter Haven, Polk County, Florida, and (B) an orchard in Windermere, Orange County, Florida. Grove A had 3 year-old Hamlin oranges on Swingle citromelo rootstocks; irrigation was applied three times per week by microjets, using reclaimed water with a high Ca and Bo content, and fertigation once a week with 9-0-9-2 NPKMg. Psyllids were controlled only occasionally. Location B had 5 year-old Hamlin and Navel oranges on Swingle citromelo rootstocks; irrigation was applied three times per week by microjets using well water, fertilizer (NPK 15-2-15) was applied once a year at about 200 kg/ha, and insecticides were applied once a month to control psyllids and mites. Nutritional sprays with a full range of macro- and micronutrients were applied twice a year at both locations. Most trees in both groves showed typical symptoms of HLB and had tested positive for Las with regular and quantitative PCR at the Division of Plant Industry, Gainesville, Florida.

Soil samples were collected on four sides under the canopy of five HLB-positive trees in each of the two groves. The surface soil (10 cm deep) was removed. Two buckets of 20 liters were filled with a spade, 20 cm deep, at the edge of the canopy. All roots with a diameter of 5 mm or less were included in each soil sample. Soil A was a yellow-brown fine sandy loam, and soil B a grey-black sandy loam. Air-dried soil samples were subjected to chemical analysis in the Soil Analysis lab at the University of Florida (UF), Gainesville, FL. Soil pH was determined with a glass electrode (soil:water = 1:5). Soil organic carbon and nitrogen, total organic matter, and soluble P and K were determined as described previously (Mylavarapu et al. 2014; Shen et al. 2013a). The pH, soluble P and K contents were very similar at the two locations, but the organic matter and total N contents were significantly higher in soil B than soil A (Table S1).

All soil was sieved through a 1-cm sieve one day after collection. The roots were cut into pieces of 1-2 cm long and returned to the soil. All tools were disinfected with 70% alcohol and all activities were carried out with clean plastic gloves to avoid cross contamination among soil samples. Next, the field capacity of each soil was determined by adding 25 ml soil and 15 ml water into funnels with Whatman 1 filter paper, draining for 30 minutes, and determining the wet weight and dry weight after drying the samples for 24 hr at 105C. Field capacities of soil A and B were 20.9% ± 1.7% and 25.5% ± 1.0%, respectively. The moisture contents of the original soil samples were 3.7% ± 0.4% and 8.4% ± 0.7% for soil A and B, respectively.

Half of the soil samples was autoclaved at 120°C in double autoclave bags for 50 min, and left open on a greenhouse bench. In order to promote colonization of the autoclaved soils by bacteria and avoid ammonia toxicity, a bacterial suspension was added, prepared from 5 g soil from an organically managed experimental vegetable field in Gainesville, mixed in 50 ml demineralized water plus 1 µl Tween 80 by vortexing, shaking for 24 hr on a rotary shaker, and sonicating in a Branson 5200 for 5 min. The suspension was centrifuged at 3400 rpm for 15 min, and filtered through a 1.2 micron sterile filter to remove fungi and protozoa. The bacterial density was determined in a spectrophotometer at 630 nm using a standard density curve after dilution plating a subsample on S-medium (O’Brien and van Bruggen 1991; van Bruggen et al. 1990). After 10-fold dilution, 200 ml of suspension was added to each subsample of 4 liters of autoclaved and cooled soil, resulting in a bacterial concentration of 10^7^ CFU/g of dry soil. The amended soil samples were mixed thoroughly in plastic bags. The microbial community was allowed to grow and equilibrate for two weeks.

#### Experimental set up and plant management

Five-month old mandarin seedlings ‘Cleopatra’ on their own roots were obtained from a citrus nursery producing certified HLB-free trees (Brite Leaf Nursery LLC, Lake Panasoffkee, Florida). The seedlings were transplanted in the autoclaved and non-autoclaved soil samples in five randomized complete blocks, one seedling per soil sample per block (for a total of 5×5×2×2=100 mandarin trees). Thirty residual seedlings were left in pasteurized potting mix. The pot size was 2 L. The pots with autoclaved and non-autoclaved soil from the same trees were placed side-by-side on greenhouse benches for paired comparisons (Fig. 1). Maintenance of the mandarin seedlings is described in the supplement.

The potted seedlings were placed on saucers to prevent potential movement of Las with irrigation water onto the ground (Florida Department of Agriculture and Consumer Services, Division of Plant Industry permit number 2009-022). The plants were watered by drip irrigation, on average 150 mL water per plant, for 1 min. every other day. The water output per day varied considerably among pots (150 ± 41 ml per watering period), but less among treatments (ranging from 126 ± 27 ml to 161 ± 21 ml per period). They were fertilized with an N-P-K (15-10-15) nutrient solution (JR Peters, Allentown, PA) with 150 mg/L of N twice a week. Once a month they also received slow release fertilizer (Osmocote 14-14-14; The Scotts Company, Marysville, OH) at 10 ml per pot. No pesticides were applied as insect pests were not observed.

The temperature in the greenhouse fluctuated between 28-33 °C during the day and 15-20 °C at night. The relative humidity fluctuated between 60 and 95%. No artificial light was provided. In the summer time, the greenhouse was shaded to reduce the daytime temperature.

#### Plant observations and sample collection

The plants were observed once a week for symptom development. Two young (but completely developed) leaves were harvested from plants with mild discoloration two months after planting using gloves sprayed with 70% alcohol before each new plant, and stored in plastic bags in the refrigerator. Two weeks later, all citrus trees in nonautoclaved soil of blocks 1 and 5, and two trees in autoclaved soil of the same blocks were transported to the laboratory. The roots were cleaned under running tapwater and inspected for insects and disease symptoms. The shoots were cut off and placed in a plastic bag, again using gloves sprayed with 70% alcohol before each new plant. Over a period of one week, petioles and midribs of all leaves stored in the refrigerator were dissected out with a sterile scalpel, weighed (107 ± 21 mg per sample), placed in eppendorf tubes and stored in the −80 freezer.

Eight and a half months after planting, the remaining citrus trees were collected as described for the sampling after two and a half months. The roots were inspected for insects and disease symptoms. The shoots were placed in a plastic bag (again, using gloves sprayed with 70% alcohol before each new plant), and stored in the refrigerator. The petioles and midribs of all leaves were dissected out, weighed, placed in eppendorf tubes and stored in the −80 freezer for DNA extraction later.

Typical symptoms of limited Pythium infection were observed on most plants in nonautoclaved soil, a common phenomenon on young plants in potted field soil. Insects were not observed during the experiment, but one plant grown in soil A, not in soil B, had citrus mealybugs (*Planococcus citri* Risso) on its foliage at the time of cleaning out the greenhouse, nine months after the start of the experiment.

---

**Table S1.** Soil chemical characteristics (pH, soluble phosphorous and potassium, total Kjeldahl nitrogen and organic matter content) of soil samples collected from two groves in central Florida in November 2009.

|  | Grove A |  | Grove B |  |
| --- | --- | --- | --- | --- |
|  | Mean | Stderr | Mean | Stderr |
| pH | 6.33 | 0.09 | 6.19 | 0.34 |
| P (mg/kg) | 2.50 | 0.07 | 4.28 | 0.19 |
| K (mg/kg) | 11.91 | 0.05 | 11.95 | 0.89 |
| TKN (mg/kg) | 337.12 | 4.35 | 800.02 | 25.16 |
| OM (%) | 0.67 | 0.01 | 1.59 | 0.05 |

P = phosphorous
K = potassium
TKN = total Kjeldahl nitrogen
OM = organic matter content

**Table S2.** *Candidatus* Liberibacter species and strains compared with *Ca*. Liberibacter terrae strain UFEPI.

| *Candidatus*<br>Liberibacter | ACC | Source | Location | Sequence<br>submission | Submission<br>year | Publication |
| --- | --- | --- | --- | --- | --- | --- |
| africanus | CP004021 | *Diaphorina citri* | South Africa | <sup>a</sup> | <sup>a</sup> | Lin et al. 2015 |
| africanus | EU921619 | Citrus | South Africa |  |  | unpublished |
| africanus | EU921620.1 | Citrus | South Africa |  |  | Lin et al. 2013 |
| africanus | KX990287.1 | *Zanthoxylum*<br>*capense* | South Africa |  |  | Roberts and Pietersen<br>2017 |
| africanus | L22533 | Periwinkle | South Africa |  |  | Jagoueix et al. 1994 |
| americanus | AY742824 | Citrus trees | Sao Paulo State<br>Brazil |  |  | Do Carmo Teixeira et<br>al. 2005 |
| americanus | EU754742.1 | Citrus | Brazil |  |  | Doddapaneni et al.<br>2008 |
| americanus | EU921624 | Citrus | Brazil |  |  | Lin et al. 2009 |
| americanus | EU999028.1 | Citrus | Brazil |  |  | Sechler et al. 2009 |
| americanus | FJ263691 | Citrus tree | Brazil |  |  | Zhou et al. 2008 |
| asiaticus | AB555706.1 | Citrus | East Timor |  |  | Miyata et al. 2011 |
| asiaticus | AP014595.1 | Citrus | Japan |  |  | Katoh et al. 2014 |
| asiaticus str. psy<br>62 | CP001677 | *Diaphorina citri* | Florida |  |  | Duan et al. 2009 |
| asiaticus | CP004005.1 | *Diaphorina citri* | China |  |  | Lin et al. 2013 |
| asiaticus | CP010804.1 | Citrus | China |  |  | Zheng et al. 2014 |
| asiaticus | CP019958.1 | Citrus | China |  |  | Zheng et al. 2017 |
| asiaticus | DQ302750.1 | Citrus | Taiwan | Tsai, C.H. et al. | 2005 | Unpublished |
| asiaticus | EU130554.1 | Citrus | USA |  |  | Keremane et al. 2008 |
| asiaticus | EU224394.1 | Citrus | Malaysia | Ahmad, K. et al. | 2007 | Publ. In Adkaretal et al. 2009 |
| asiaticus | EU644449 | *Citrus reticulata* | China |  |  | Deng et al. 2008 |
| asiaticus | EU921615 | Citrus | China |  |  | Lin et al. 2009 |
| asiaticus | EU921616 | Citrus | China |  |  | Lin et al. 2009 |
| asiaticus | EU921622.1 | Citrus | Brazil |  |  | Lin et al. 2009 |
| asiaticus | EU921617 | Citrus | China |  |  | Lin et al. 2009 |
| asiaticus | FJ196314 | Kinnow mandarin | India |  |  | Gupta et al. 2012 |
| asiaticus | FJ750456 | Sweet orange | Florida |  |  | Unpublished |
| asiaticus | FJ750459 | Satsuma plant | Lousiana |  |  | Unpublished |
| asiaticus | JQ867409.1 | Citrus | Mexico | Moreno, A. et al. | 2012 | Unpublished |
| asiaticus | JQ867411.1 | Citrus | Mexico | Moreno, A. et al. | 2012 | Unpublished |
| asiaticus | JQ867413.1 | Citrus | Mexico | Moreno, A. et al. | 2012 | Unpublished |
| asiaticus | JQ867424.1 | Citrus | Mexico | Moreno, A. et al. | 2012 | Unpublished |
| asiaticus | JQ867452.1 | Citrus | Mexico | Moreno, A. et al. | 2012 | Unpublished |
| asiaticus | KC551939.1 | Citrus | India | Das, AK et al. | 2013 | Unpublished |
| asiaticus | KU761591.1 | Citrus sp., Kagzi lime | India | Chattopadhiyay, D. et al. | 2016 | Unpublished |
| asiaticus | KX218368.1 | Citrus sp., Kagzi lime | India | Chattopadhiyay, D. and Sarkar, K. | 2016 | Unpublished |
| asiaticus | KY323722.1 | Citrus | Iran |  |  | Alizadeh et al. 2017 |
| asiaticus | KY990822.1 | Citrus sp., Kagzi lime | India | Chattopadhiyay, D. et al. | 2016 | Unpublished |
| asiaticus | L22532 | Periwinkle | India |  |  | Jagoueix et al. 1994 |
| asiaticus | MH368772.1 | Citrus | China | Munir, S., and He, Y. | 2018 | Unpublished |
| asiaticus | MH368774 | Citrus | China | Munir, S., and He, Y. | 2018 | Unpublished |
| crescens | KY604742.1 | *Muraya koenigii* | USA | Rascoe, J. et al. | 2018 | Unpublished |
| psyllaorous | EU812557 | *Bactericera cockerelli* | California |  |  | Hansen et al. 2008 |
| solanacearum | EU935004 | *Solanum*<br>*betaceum* | New Zealand |  |  | Liefting et al. 2009 |
| solanacearum | MG701017.1 | Trioza anthrisci | Finland |  |  | Haapalainen et al. 2018 |
| Sp. | AB008366.1 | Citrus | Japan | Sabandiyah, S. et al. | 2007 | Unpublished |
| Sp. | LN835770.1 | Citrus | Pakistan |  |  | Zafarullah et al. 2016 |
| terrae strain<br>UFEPI | MK125061 | Mandarin | Florida |  |  | This study |
| Uncultured<br>bacterium M-20 | GU553033.1 | *Diaphorina citri* | China | Tian, S.C. et al. | 2010 | Unpublished |

<sup>a</sup> Sequences not found in Genbank

**Table S3.** Ct values with primers specific to *Candidatus* Liberibacter asiaticus (Li et al. 2006) in quantitative real-time TaqMan PCR assays in mandarin midribs plant extracts.

| Plant Sample designations<sup>a</sup> | Plant sample numbers submitted to Genbank | Accession number<sup>b</sup> | Soil location | Sampling time (months) | Ct value |
| --- | --- | --- | --- | --- | --- |
| 1AR1T9 | CaLUF1 | MF463003 | B | 2.5 | 31.8 |
| 1AR4T1 | CaLUF2 | MF463004 | B | 2.5 | 32.9 |
| 2AR2T2 | CaLUF3 | MF463005 | B | 2.5 | 31.7 |
| 4AR4T1 | CaLUF4 | MF463006 | B | 2.5 | 32.0 |
| 5AR1T9 | CaLUF5 | MF463007 | B | 2.5 | 29.5 |
| 2BR2T2 | CaLUF6 | MF463008 | B | 8.5 | 33.9 |
| 4BR4T1 | CaLUF7 | MF463009 | B | 8.5 | 33.8 |

<sup>a</sup> The first number designates the plant number in the greenhouse, followed by the sampling time (A or B); the third and fourth numbers indicate the row and tree numbers in the citrus grove where soil had been collected.
<sup>b</sup> The accession number obtained from Genbank

**Table S4.** 16S rDNA gene pairwise similarity among *Candidatus* Liberibacter terrae str. UFEPI obtained from 7 mandarin plants grown in soil from an HLB-infested citrus grove.

| # | *Candidatus*<br>Liberibacter<br>terrae strain<br>UFEPI | 1 | 2 | 3 | 4 | 5 | 6 |
| --- | --- | --- | --- | --- | --- | --- | --- |
| 1 | 1AR1T9 |  |  |  |  |  |  |
| 2 | 1AR4T1 | 0.000000 |  |  |  |  |  |
| 3 | 2AR2T2 | 0.000955 | 0.000955 |  |  |  |  |
| 4 | 4AR4T1 | 0.000955 | 0.000955 | 0.001909 |  |  |  |
| 5 | 5AR1T9 | 0.000955 | 0.000955 | 0.001909 | 0.001909 |  |  |
| 6 | 2BR2T2 | 0.000000 | 0.000000 | 0.000955 | 0.000955 | 0.000955 |  |
| 7 | 4BR4T1 | 0.000000 | 0.000000 | 0.000955 | 0.000955 | 0.000955 | 0.000000 |

**Table S5.** Pairwise similarity of the β-operon gene among *Candidatus* Liberibacter terrae strains UFEPI obtained from 7 mandarin plants grown in soil from an HLB-infested citrus grove. Note that all the isolates are more than 99.6% similar.

| # | *Candidatus*<br>Liberibacter<br>terrae strains | 1 | 2 | 3 | 4 | 5 | 6 |
| --- | --- | --- | --- | --- | --- | --- | --- |
| 1 | 1AR1T9 |  |  |  |  |  |  |
| 2 | 1AR4T1 | 0.000432 |  |  |  |  |  |
| 3 | 2AR2T2 | 0.001765 | 0.000955 |  |  |  |  |
| 4 | 4AR4T1 | 0.003915 | 0.000955 | 0.001909 |  |  |  |
| 5 | 5AR1T9 | 0.003255 | 0.000955 | 0.001909 | 0.001909 |  |  |
| 6 | 2BR2T2 | 0.000310 | 0.000000 | 0.000955 | 0.000955 | 0.000955 |  |
| 7 | 4BR4T1 | 0.000000 | 0.000000 | 0.000955 | 0.000955 | 0.000955 | 0.000000 |

**Fig. S1.**
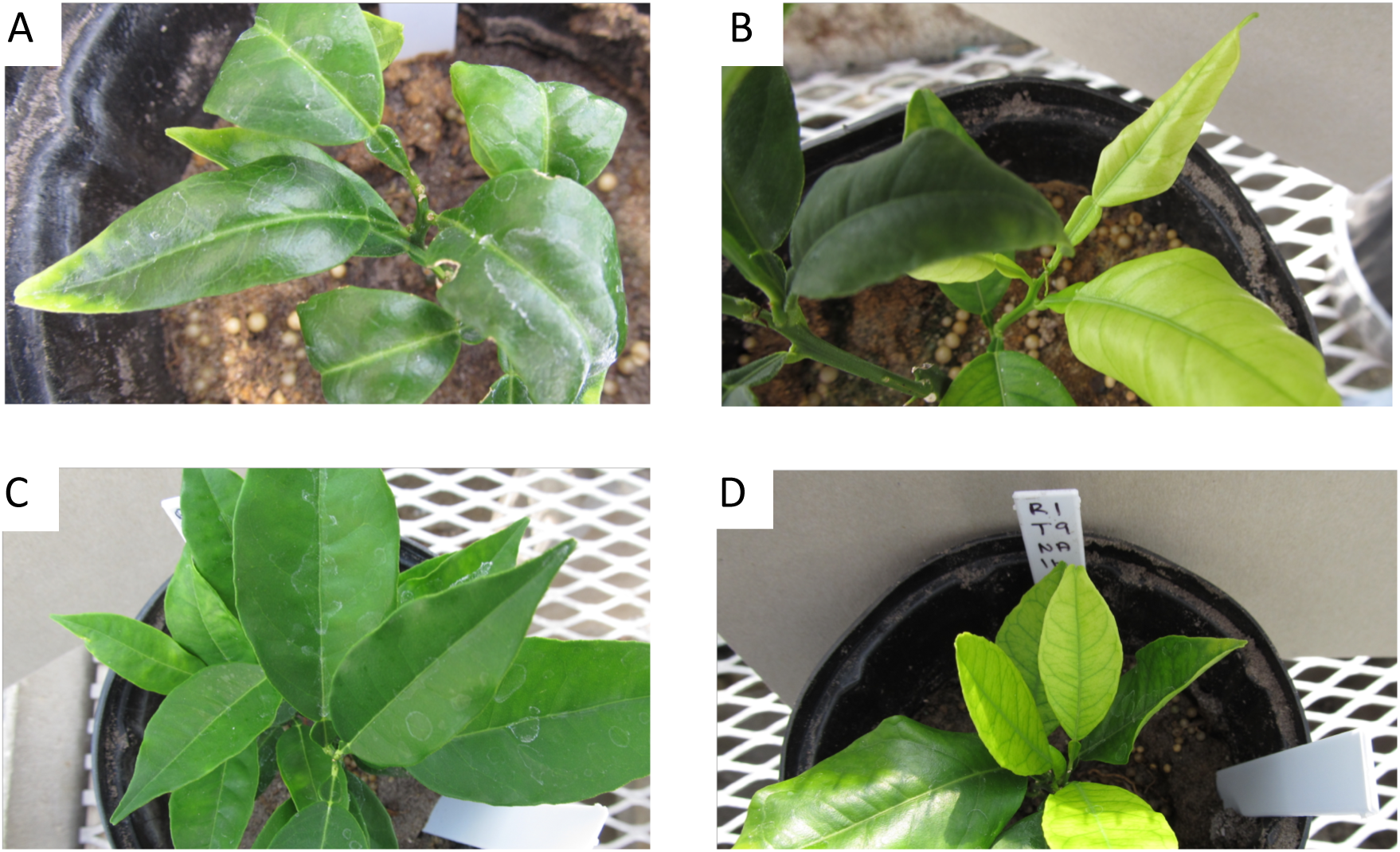
Mandarin seedlings four months after planting in autoclaved (A and C) and non-autoclaved (B and D) soil from under the canopy of Hamlin trees with typical huanglongbing symptoms in grove B. Soil samples in A and B were collected under tree R2T2 and soil samples in C and D under tree R1T9. DNA extracts from seedlings in non-autoclaved soil samples R2T2 and R1T9 (B and D) contained the new Ca. Liberibacter strain UFEPI 2.5 months after planting.

**Fig. S2.**
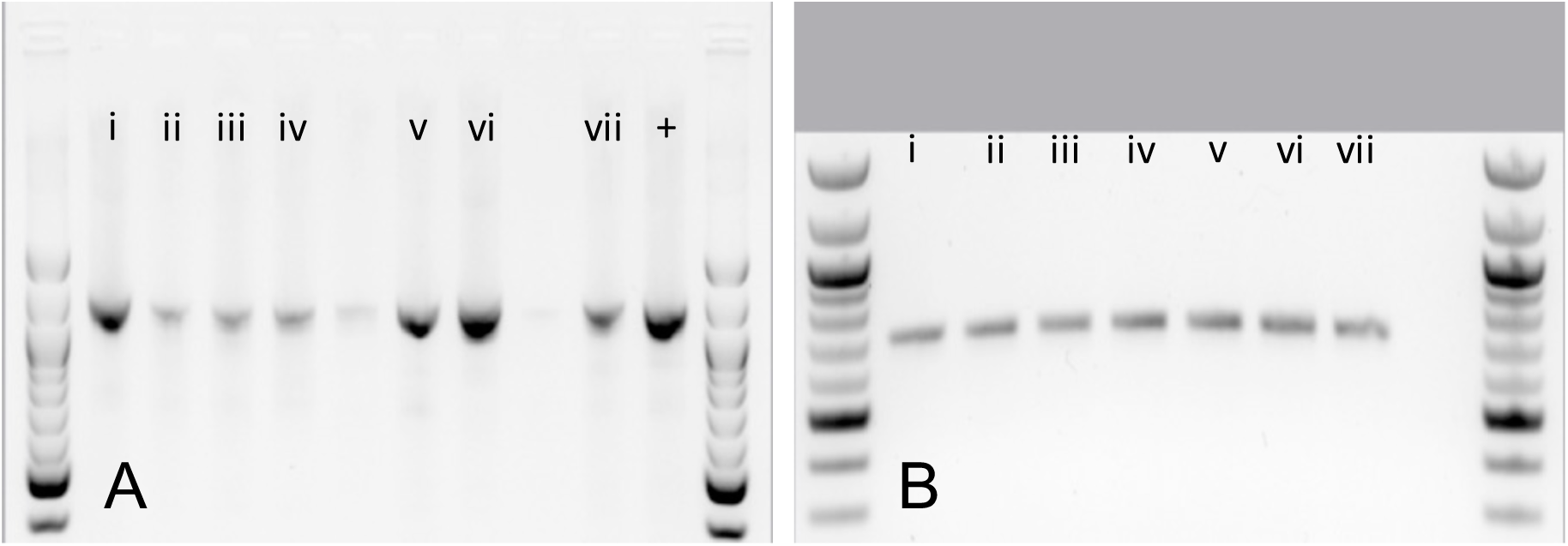
PCR products for (A) *Candidatus* Liberibacter spp. 16S rDNA gene group-specific primers and (B) *Candidatus* Liberibacter spp. β-operon group specific primers. (i) 1AR1T9, (ii) 1AR4T1, (iii) 2AR2T2, (iv) 4AR4T1, (v) 5AR1T9, (vi) 2BR2T2 and (vii) 4BR4T1, (+) positive *Candidatus* Liberibacter asiaticus sample.

**Fig. S3.**
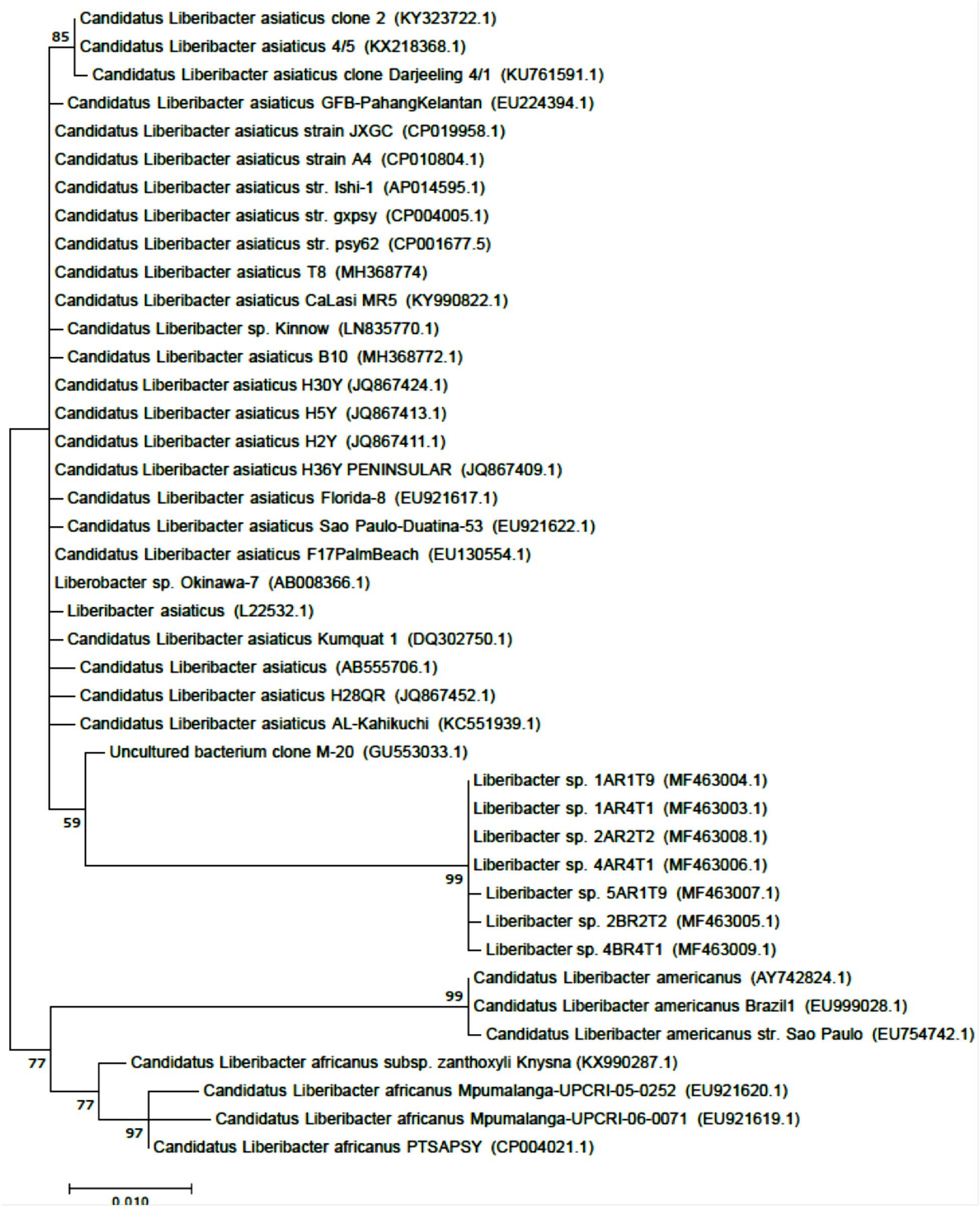
Phylogenetic tree based on maximum-likelihood analysis of the 16S rDNA sequences with various members within the genus *Candidatus* Liberibacter without outgroup. Phylogeny was inferred using the Tamura-Nei model (Tamura and Nei 1993) with gamma correction to account for site variations. Bootstrap values based on 1,000 replications are shown at branch nodes and values under 50 are not shown. Genbank accessions are shown on the tree for sequences indicated in brackets. Bar, 0.010 substitutions per nucleotide position.

**Fig. S4.**
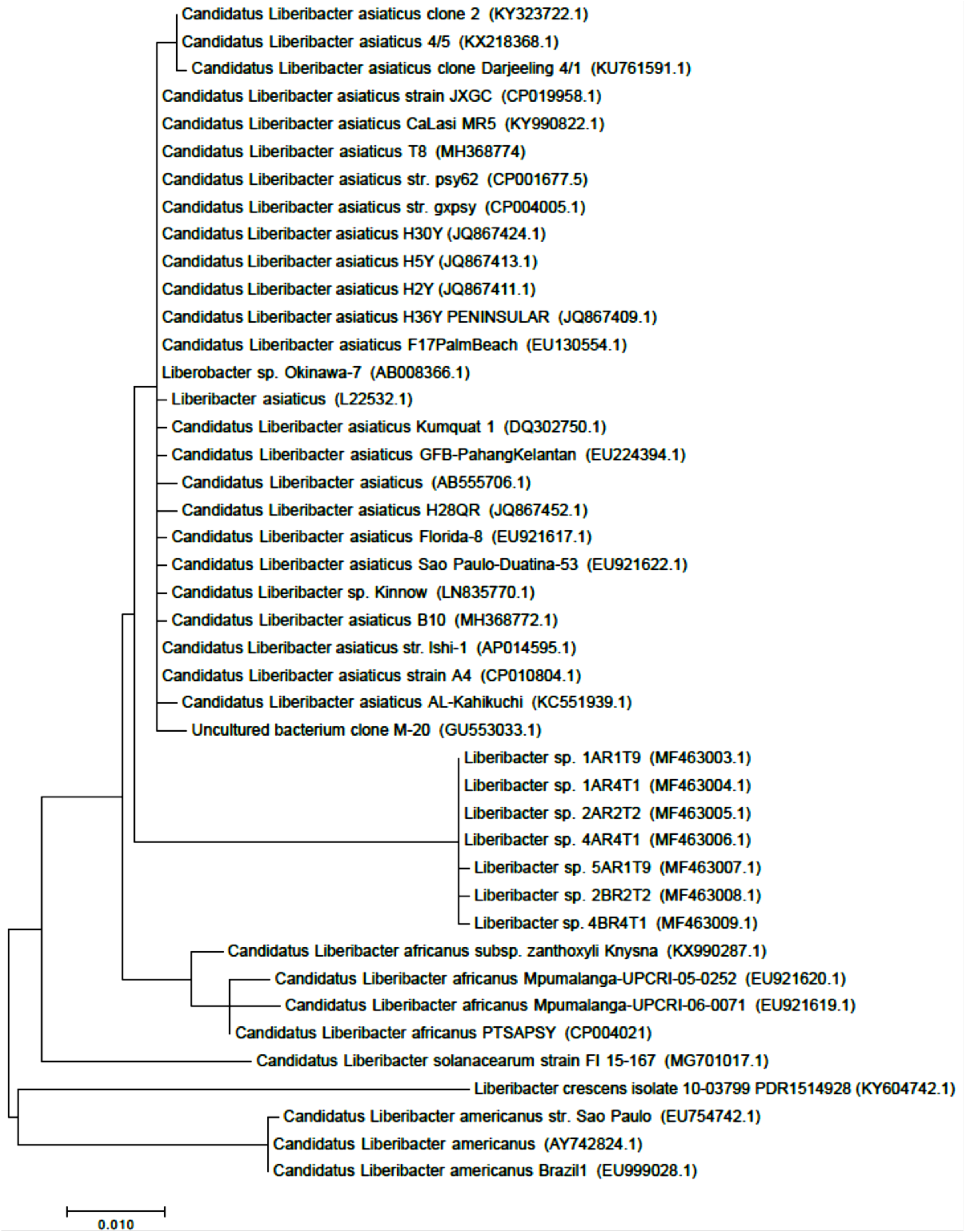
Phylogenetic tree based on maximum-likelihood analysis of the 16S rDNA sequences with various members within the genus *Candidatus* Liberibacter and *Liberibacter crescens* without outgroup. Phylogeny was inferred using the Tamura-Nei model (Tamura and Nei 1993) with gamma correction to account for site variations. Genbank accessions are shown on the tree for sequences indicated in brackets. Bar, 0.010 substitutions per nucleotide position.

